# Behavioral thermal tolerance predicts distribution pattern but not habitat use in sympatric Neotropical frogs

**DOI:** 10.1101/2020.04.04.024612

**Authors:** Juan C. Diaz-Ricaurte, Filipe C. Serrano, Estefany Caroline Guevara-Molina, Cybele Araujo, Marcio Martins

## Abstract

Environmental temperatures are a major constraint on ectotherm abundance and diversity, influencing their distribution and natural history. Comparing thermal tolerances with environmental temperatures is a simple way to estimate thermal constraints on species distributions. We investigate the potential effects of thermal tolerance on anuran local (habitat) and global distribution patterns and associated behavioral responses. We tested for differences in Voluntary Thermal Maximum of two sympatric frog species of the genus *Physalaemus* in the Cerrado ecoregion. For each species, we constructed models to assess the effects of period of day, duration of experiment, initial body mass, initial body temperature and heating rate on the VT_Max_. We mapped the difference between VT_Max_ and maximum daily temperature (VT_Max_ - ET_Max_) for each occurrence point. *Physalaemus nattereri* had a significantly higher VT_Max_ than *P. cuvieri*. For *P. nattereri*, the model including only period of day was chosen as the best to explain variation in the VT_Max_. For *P. cuvieri*, no model was selected as best to predict VT_Max_. The VT_Max_ - ET_Max_ values were significantly different between species, with *P. nattereri* mostly found in localities that attain maximum temperatures lower than its VT_Max_ and *P. cuvieri* showing the reverse pattern. Regarding habitat use, we found *P. cuvieri* to be slightly more abundant in open habitats than in non-open habitats, whereas *P. nattereri* shows the reverse pattern. The difference in VT_Max_ values between these two species might be related to their different body sizes, but additionally might reflect their natural history, especially the way they use their habitats, and phylogenetic constraints (the species studied are in different clades within *Physalaemus*). Our study indicates that differences in behavioral thermal tolerance may be important in shaping local and regional distribution patterns. Furthermore, small-scale habitat use might reveal a link between behavioral thermal tolerance and natural history strategies.

## Introduction

Environmental temperatures are a major constraint on ectotherm abundance and diversity, influencing their distribution and natural history [1, 2, 3]. Several studies have explored environmental constraints on ectothermic vertebrates at regional and global scale [4, 5, 6]. The physiological performance of individuals can be negatively affected by high environmental temperatures [7], which can lead to declining populations and/or local extinctions [3]. Thus, knowing species thermal tolerance and exploring how environmental temperatures might affect their physiology and restrict their distribution is of primary concern for long-term conservation, especially under a global warming trend (e.g. [8]).

Many studies infer potential distribution of species using solely environmental temperatures from occurrence localities to model their potential niche (e.g. [9–10]). However, incorporating thermal tolerances to these analyses allows a more realistic approach to estimate thermal constraints on distributions [11]. Behavioral and physiological thermal tolerances impact not only species ranges, but also the distribution and abundance patterns of their populations [3]. Identifying thermal tolerance thresholds outside the range of preferential body temperatures for thermoregulation (see [12]) allows for the identification of temperatures that directly affect the behavioral and physiological thermal tolerance of ectothermic organisms. One of the thresholds related to PBT is the Voluntary Thermal Maximum (VT_Max_), which represents a behavioral thermal tolerance, i.e., the maximum temperature that an organism will endure before trying to move to a place with a lower temperature, thus trying to maintain its body temperature within its range of preferential body temperatures [3, 12, 13]. However, if an individual fails to respond to its VT_Max_, an increase in body temperature will expose it to its physiological thermal limit (i. e., its Critical Thermal Maximum), which can lead to functional collapse and consequently death due to overheating [13, 14].

Behavioral thermal tolerances can be influenced by factors such as reproductive status, sex, photoperiod, and hydration state [12, 15]. Additionally, thermal tolerances such as the VT_Max_ might decrease with body size: due to thermal inertia, larger animals might have slower heating and cooling rates than small animals, which increases the exposed time to stressful thermal conditions [16–17]. Yet, most studies focus only on upper and critical temperatures (e.g. [18–19] and thus fail to incorporate behavioral response, which derives from the animal’s own perception of thermal stress. Additionally, the behavioral response to upper voluntary limits might represent a more informative ecological threshold to identify thermal constraints on geographic distribution [3, 8, 13, 20–25]. Contrary to the Critical Thermal Maximum, the exposure to the VT_Max_ does not induce an immediate loss of locomotion. Thus, VT_Max_ can more realistically portrait changes in species behavior associated with their natural history.

The genus *Physalaemus* Fitzinger 1826 is one of the largest groups of frogs in the Neotropics, with 48 recognized species [26]. This group is distributed from the lowlands of southern Venezuela and the Llanos of southeastern Colombia to central Argentina [26]. Some species of this genus have sympatric populations along extensive areas, such as *Physalaemus nattereri* [27] and *Physalaemus cuvieri* [28] (see [26]), which are widespread in central South America [29]. These species belong to different clades within *Physalaemus* (*P. signifer* and *P. cuvieri* clades, respectively; [30]. *Physalaemus nattereri* has a stout body, a moderate to large size (adult snout-to-vent length of 29.8–50.6 mm) and is endemic to the Cerrado, whereas *P. cuvieri* has a slenderer body, a small size (snout-to-vent length of adults 28–30 mm) and occurs throughout the Cerrado, in southern portions of the Amazon Forest and in the Atlantic Forest [31]; but see Methods). Even though both species occur in Cerrado, it is still unclear how their local abundances vary within Cerrado vegetation types, from the ‘cerradão’, which has a forest structure, to grasslands (‘campo sujo’). Additionally, *P. cuvieri* and *P. nattereri* differ in their biology. While *P. cuvieri* uses several aquatic habitats for reproduction and seeks shelter during the day in already-dug burrows, *P. nattereri* breeds mostly in temporary puddles and buries itself in the soil during the day aided by metatarsal tubercles (S1 Fig) on its hind feet [32–34].

Herein we investigate the potential effects of temperature on anuran local (habitat) and global distribution patterns and associated behavioral responses. We test for differences in the VT_Max_ between *P. nattereri* and *P. cuvieri* and explore possible effects of their behavioral thermal tolerances on their geographic distribution and habitat use in the Cerrado. We hypothesize that: i) the VT_Max_ is lower for *P. nattereri* due to its larger body size and consequent slower cooling rates; ii) the species with the lower VT_Max_ is less abundant in habitats with higher environmental temperatures, such as open areas of the Cerrado; and iii) there is a relationship between VT_Max_ and geographic distribution, such that species occur mostly in localities where the maximum environmental temperature is below their VT_Max_. We expect that our results can contribute to studies on the effect of global climate change on habitat use and geographic distribution.

## Materials and Methods

### Physiological Parameters

#### Capture and maintenance of individuals

Fieldwork was carried out at Estação Ecológica de Santa Bárbara (22°49’2.43”S, 49°14’11.29”W; WGS84, 590 m elevation), one of the few remnants of Cerrado savannas in the state of São Paulo, Brazil, with a total area of 2,712 ha [35]. The climate is Humid subtropical [36], with temperatures averaging 24°C and 16°C during January and July, the hottest and coldest months, respectively. The average annual rainfall is 1100–1300 mm, with marked dry and wet seasons (approximately April to September and October to March, respectively; [37]). The landscape consists mainly of open grassland and savanna-type formations, such as ‘campo sujo’ and ‘campo cerrado’, but also of non-open vegetation types such as ‘cerrado *strictu sensu*’ (dense savanna) and ‘cerradão’ (cerrado woodland). Between 24 and 28 September 2018, we captured 14 individuals of *P. nattereri* and 20 of *P. cuvieri* in pitfall traps with drift fences [38–39] and these individuals were housed individually in plastic boxes at room temperature.

#### Measurements of the Voluntary Thermal Maximum (VT_Max_)

To obtain the VT_Max_ for each species, we measured each individual within one day of collection at 100% hydration level. To reach maximum hydration level, each individual was placed in a cup with water *ad libitum* one hour prior to the experiment. Then its pelvic waist was pressed to expel the urine and to obtain its 100% hydration level in relation to its standard body mass. We heated each individual in a metal box wrapped in a thermal resistance for heating. The box had a movable lid, allowing the animal to easily leave the box when needed. A thin thermocouple (type-T, Omega®) was located in the inguinal region of each individual to record its body temperature during the heating experiment [15]. Another type-T thermocouple was placed on the surface of the box to record heating rate and to make sure that the temperature did not exceeded 5–6°C the temperature of the individual. The thermocouples were calibrated and connected to a FieldLogger PicoLog TC-08 to record temperature data every 10 seconds. The VT_Max_ of each individual was recorded as its last body temperature at the time of leaving the box. Once its final body mass was measured, it was taken to a container with water for recovery. Furthermore, to control for a potential circadian effect on tolerances, we tested if the VT_Max_ differed between different times of the day by testing half of the individuals of each species in different periods: 10h00 to 17h00 (daytime) and 19h00 to 24h00 (nighttime).

#### Statistical analyses

We used t-tests to compare the VT_Max_, period (day or night), duration of experiment, initial body mass, initial body temperature and heating rate between species. For each species, we constructed generalized least squares models to assess the effects of the previous variables on the VT_Max_. We used the Akaike Information Criterion (AIC) to select the model that best represented the effects of factors and their interaction on the VT_Max_ of each species. Differences of two units in AIC were not considered to be different [40]. We considered the model with AIC weight values close or equal to 1 to represent the strongest model. All statistical analyses and plotting were performed in R 3.5.0 [41], with the nlme [42], ggplot2 [43] and AICcmodavg [44] packages.

#### Distribution and habitat

We used vouchered occurrence data for *P. cuvieri* (N = 163) and *P. nattereri* (N = 164) in the Cerrado from a distribution database built for another study [45]. Although the populations traditionally assigned to *P. cuvieri* (see [26]) may include more than one cryptic species (see [30]), most of the distribution of *P. cuvieri* in the Cerrado correspond to a single lineage (Lineage 2 in [30]). We calculated and mapped VT_Max_ - ET_Max_ as the difference between the VT_Max_ and maximum environmental temperature (Bio 5; 30 seconds resolution from WorldClim Vr. 2.0; [46]) for each occurrence point in Cerrado and the VT_Max_ obtained at Estação Ecológica de Santa Bárbara for *P. cuvieri* and *P. nattereri*. We used a t-test to compare VT_Max_ - ET_Max_ of species ocurrence points. All maps and GIS procedures were made in QGIS 3.12 [47]. We tested for differences between species in habitat use by comparing abundances in open (‘campo cerrado’, ‘campo sujo’ and ‘campo limpo’) and non-open habitats (gallery forest, ‘cerradão’ and cerrado *stricto sensu*; [48] for communities within Cerrado where both species occur in sympatry [49–50].

## Results

### Voluntary Thermal Maximum (VT_Max_) and experimental conditions

We found that the mean VT_Max_ was significantly lower in *P. cuvieri* than in *P. nattereri* (Table 1; t = 3.99, df = 32, p = 0.0003). We also found significant differences in initial body mass (Table 1; t = 4.26, df = 32, p = 0.0001) between species. We did not find significant differences in start body temperatures (Table 1; t = 1.39, df = 32, p = 0.1735), period of day (Table 1; t = 0.12, df = 32, p = 0.9051), duration of the experiment (Table 1; t = 0.44, df = 32, p = 0.6585) and heating rate (Table 1; t = 1.51, df = 32, p = 0.1395) between species (see S1, S2 and S3 Tables).

**Table 1.**
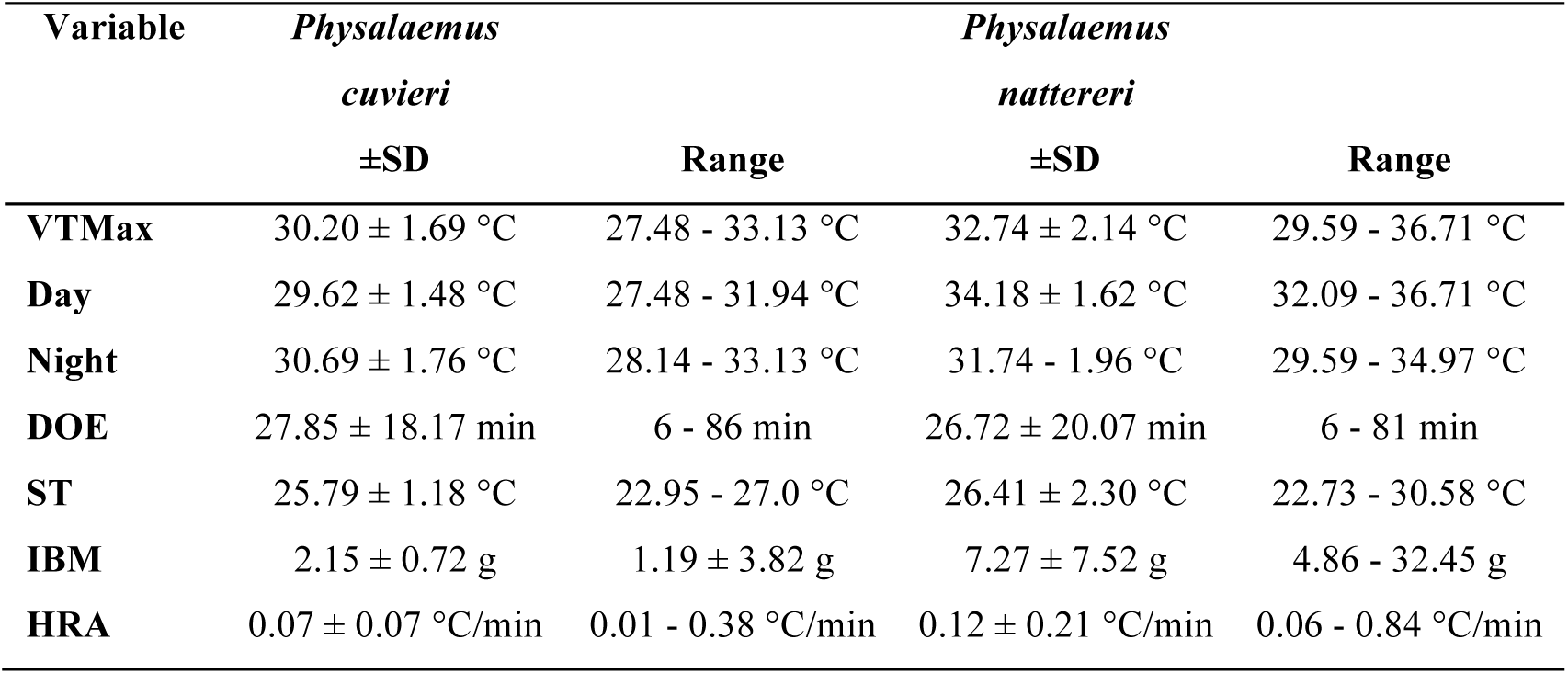
Variation of the VT_Max_ and experimental variables for *P. cuvieri* and P. *nattareri* from Estação Ecológica de Santa Bárbara, state of São Paulo, Brazil. Experimental variables are: period of day (day and night), initial body temperature (ST), duration of experiment (DOE), initial body mass (IBM), and heating rate (HRA).

We tested six models for both species using the AIC selection criteria. For *P. nattereri*, the model including only period (day or night) was chosen as a better explanation of variation in the VT_Max_ (Table 2), with higher values attained during daytime. For *P. cuvieri*, the model considering only the period of experiment had the highest AIC weight. However, the difference of its AIC value in relation to the null model was less than two units and thus we retained the simpler null model, which indicates that no variable explains the variation of the VT_Max_ of this species (Table 3).

**Table 2.** Effect of period, start body temperature, duration, initial body mass, and heating rate on the Voluntary Thermal Maximum (VT_Max_) of *P. nattereri* from Estação Ecológica de Santa Bárbara, state of São Paulo, Brazil.

| Mode<br>I | Variables | Value | Std.Error | t-value | p-value | AIC-value | AIC-weight |
| --- | --- | --- | --- | --- | --- | --- | --- |
| VI | Intercept | 34.245 | 0.746 | 45.89 | 0 | 60.46 | 0.68 |
|  | Period | -2.393 | 0.987 | -2.424 | 0.032 |  |  |
| I | Intercept | 32.877 | 0.572 | 57.384 | <0.0001 | 64.043 | 0.22 |
| V | Intercept | 33.115 | 6.588 | 5.026 | 0.0004 | 62.422 | 0.09 |
|  | Period | -2.340 | 1.075 | -2.176 | 0.0522 |  |  |
|  | Start body temperature | 0.041 | 0.239 | 0.172 | 0.8661 |  |  |
| IV | Intercept | 32.907 | 7.067 | 4.655 | 0.0009 | 64.396 | 0.01 |
|  | Period | -2.382 | 1.167 | -2.040 | 0.0686 |  |  |
|  | Start body temperature | 0.043 | 0.251 | 0.174 | 0.8652 |  |  |
|  | Duration | 0.003 | 0.028 | 0.137 | 0.8937 |  |  |
| II | Intercept | 40.452 | 8.555 | 4.728 | 0.0015 | 64.076 | 0 |
|  | Period | -4.717 | 1.818 | -2.594 | 0.0319 |  |  |
|  | Start body temperature | -0.261 | 0.308 | -0.845 | 0.4225 |  |  |
|  | Duration | 0.047 | 0.038 | 1.234 | 0.2519 |  |  |
|  | Initial body mass | 0.133 | 0.088 | 1.511 | 0.1691 |  |  |
|  | Heating rate | -5.611 | 4.139 | -1.355 | 0.2122 |  |  |
| III | Intercept | 33.394 | 7.097 | 4.704 | 0.0011 | 64.971 | 0 |
|  | Period | -2.897 | 1.281 | -2.260 | 0.0501 |  |  |
|  | Start body temperature | -0.007 | 0.257 | -0.030 | 0.9762 |  |  |
|  | Duration | 0.013 | 0.03 | 0.443 | 0.6679 |  |  |
|  | Initial body mass | 0.081 | 0.083 | 0.981 | 0.3518 |  |  |

**Table 3.** Effect of period, start body temperature, duration, initial body mass, and heating rate on the Voluntary Thermal Maximum (VT_Max_) of *P. cuvieri* from Estação Ecológica de Santa Bárbara, state of São Paulo, Brazil.

| Mode<br>I | Variables | Value | Std.Error | t-value | p-value | AIC-value | AIC weight |
| --- | --- | --- | --- | --- | --- | --- | --- |
| I | Intercept | 30.263 | 0.373 | 81.123 | <0.0001 | 80.204 | 0.44 |
| VI | Intercept | 29.623 | 0.533 | 55.479 | 0 | 79.497 | 0.42 |
|  | Period | 1.163 | 0.719 | 1.615 | 0.1236 |  |  |
| V | Intercept | 28.823 | 8.404 | 3.429 | 0.0032 | 81.486 | 0.09 |
|  | Period | 1.144 | 0.766 | 1.493 | 0.1536 |  |  |
|  | Start body temperature | 0.031 | 0.329 | 0.095 | 0.9251 |  |  |
| IV | Intercept | 24.78 | 8.621 | 2.874 | 0.011 | 81.05 | 0.05 |
|  | Period | 1.516 | 0.786 | 1.927 | 0.0719 |  |  |
|  | Start body temperature | 0.131 | 0.327 | 0.402 | 0.6925 |  |  |
|  | Duration | 0.032 | 0.022 | 1.439 | 0.1693 |  |  |
| III | Intercept | 24.608 | 8.938 | 2.753 | 0.0148 | 83.001 | 0.01 |
|  | Period | 1.574 | 0.866 | 1.817 | 0.0892 |  |  |
|  | **Start body temperature** | 0.144 | 0.343 | 0.419 | 0.6807 |  |  |
|  | **Duration** | 0.034 | 0.026 | 1.323 | 0.2054 |  |  |
|  | **Initial body mass** | -0.115 | 0.601 | -0.191 | 0.8505 |  |  |
| **II** | **Intercept** | 25.217 | 9.371 | 2.69 | 0.0176 | 84.824 | 0 |
|  | **Period** | 1.56 | 0.893 | 1.746 | 0.1027 |  |  |
|  | **Start body temperature** | 0.128 | 0.357 | 0.358 | 0.7255 |  |  |
|  | **Duration** | 0.034 | 0.026 | 1.285 | 0.2194 |  |  |
|  | **Initial body mass** | -0.082 | 0.626 | -0.131 | 0.8973 |  |  |
|  | **Heating rate** | -1.906 | 5.408 | -0.352 | 0.7297 |  |  |

### Distribution and habitat

Overall distribution of occurrences was similar for the two species, occupying mainly the central and southern portions of the Cerrado (Fig 1; S4 Table). Thus, the distribution of environmental temperatures was similar for both species. However, because the VT_Max_ was different between species, the resulting distribution of VT_Max_ - ET_Max_ values were significantly different (Fig 1A–B). The north central portion of the Cerrado showed much higher environmental temperatures than the VT_Max_ of *P. cuvieri* (Fig 1A), while this region is mostly below the VT_Max_ of *P. nattereri* (Fig 1B). Furthermore, VT_Max_ - ET_Max_ values were found to be significantly different between species (t = 13.26, df = 214.98, p < 0.001; Fig 1C). *Physalaemus nattereri* is mostly found (∼ 80%) on localities that attain maximum temperatures equal to or lower than its VT_Max_, whereas *P. cuvieri* seems to be mostly distributed (∼ 60%) in localities with higher VT_Max_ - ET_Max_ values (Fig 1C).

**Fig 1.**
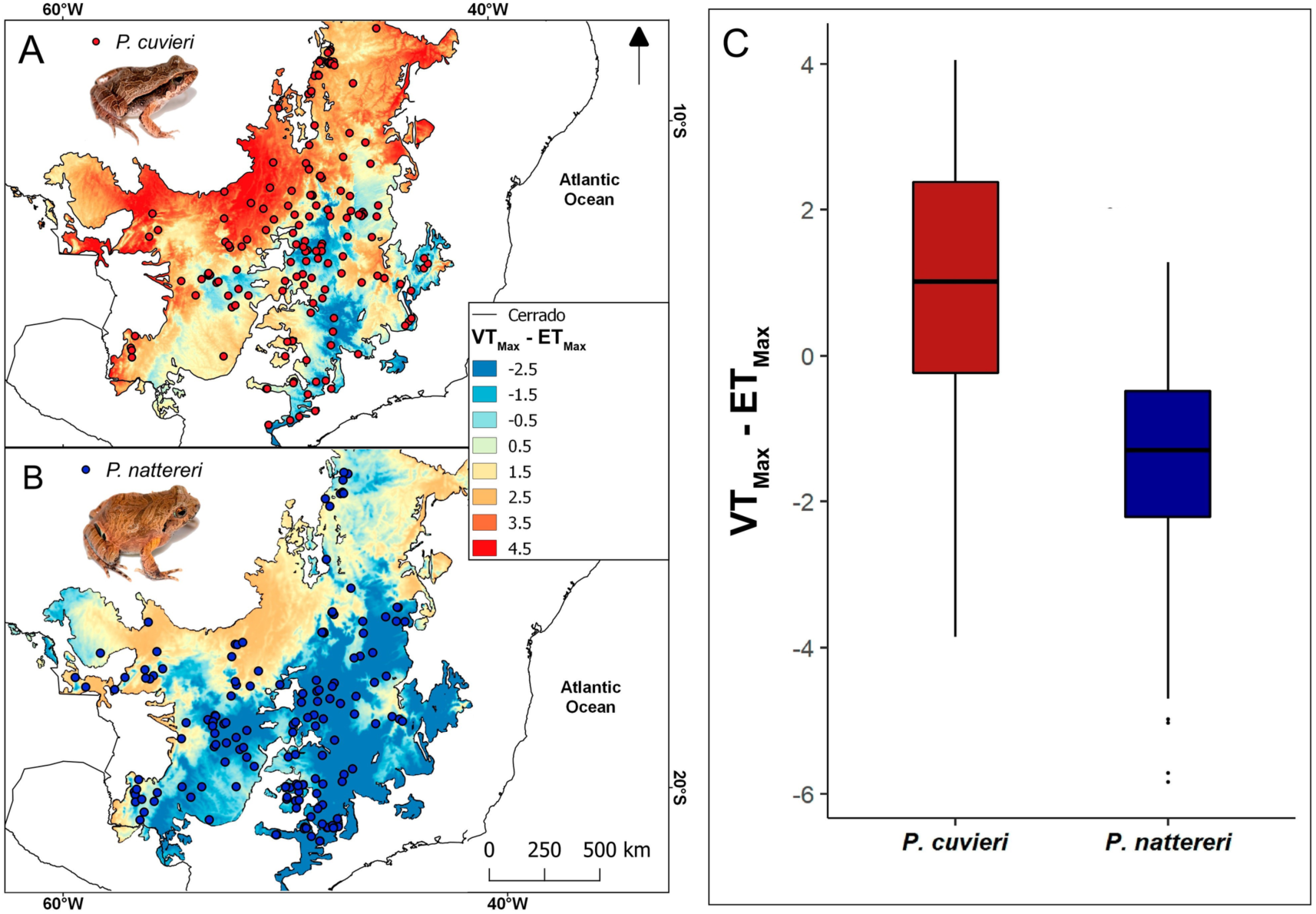
Geographical distribution of two species of frogs and VT_Max_ -ET_Max_ values throughout their distribution. (A) Distribution of *Physalaemus cuvieri*; (B) Distribution of *Physalaemus nattereri*; and C) Boxplot of VT_Max_ - ET_Max_ values at occurrence points for both species in the Cerrado.

We obtained abundance data for six localities in southern Cerrado, most of them from protected areas (Fig 2). *Physalaemus cuvieri* was more abundant in open areas than in non-open areas (mean = 58.1%; sd = 35.5%), while *P. nattereri* was less abundant in open areas than in non-open areas (mean = 42.3%; sd = 29.5%; S5 Table).

**Fig 2.**
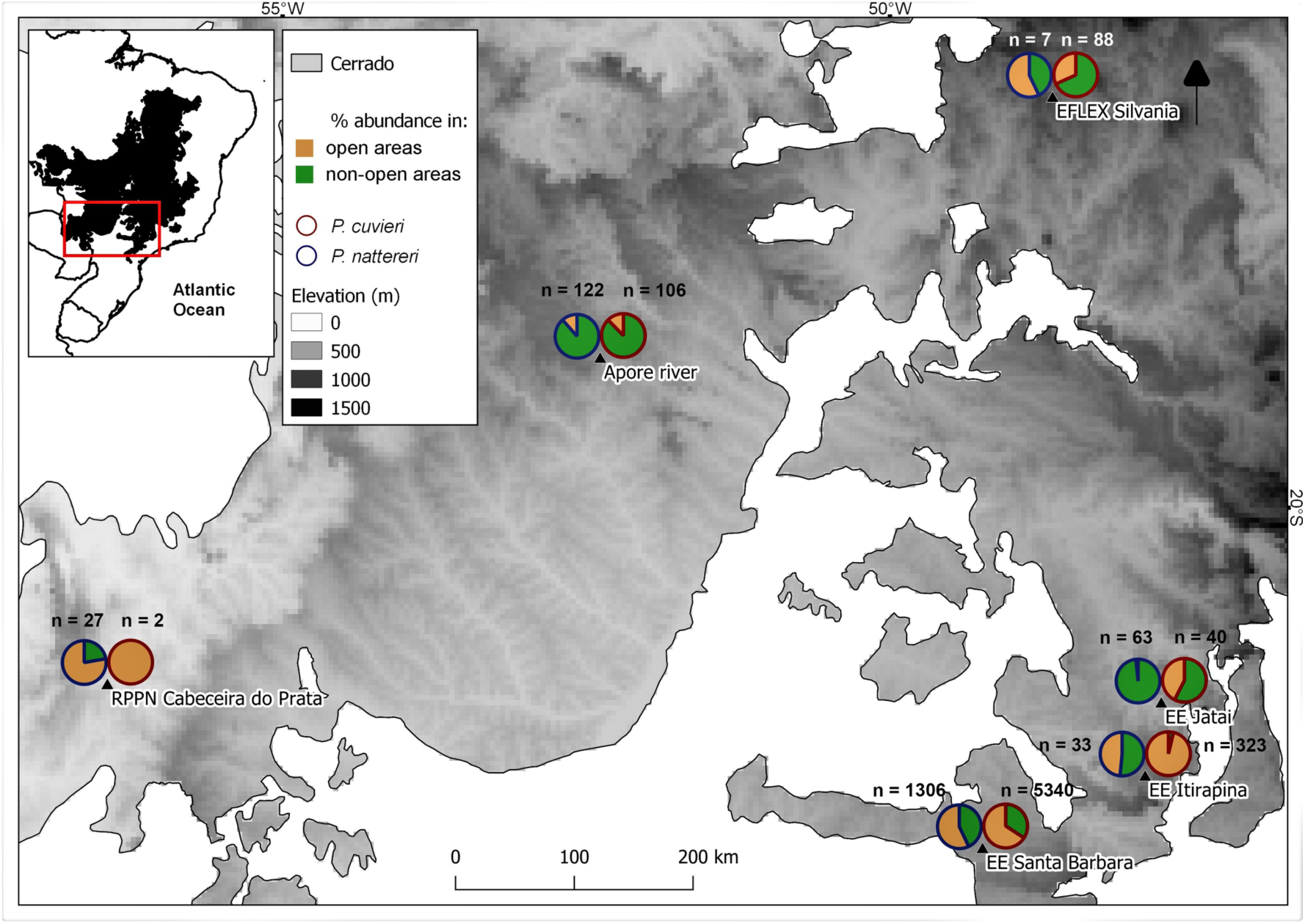
Relative abundance of *P. cuvieri* and *P. nattereri* in open and non-open areas in Cerrado. Relative abundance (in %) of *P. cuvieri* (blue circles) and *P. nattereri* (red circles) in open (brown) and non-open areas (green) in Cerrado in six localities (see Table S5): Estação Florestal de Experimentação (EFLEX) de Silvânia (GO), Reserva Particular do Patrimônio Natural (RPPN) Cabeceira do Prata (MS), Estação Ecológica (EE) Jataí (SP), Estação Ecológica de Itirapina (SP), Estação Ecológica de Santa Bárbara (SP), and Aporé river (GO and MS). Sources of data: [49, 51–54].

## Discussion

Our results show that the Voluntary Thermal Maximum (VT_Max_) is higher for *P. nattereri* than for *P. cuvieri*, contrary to our first prediction that larger body size (and an expected slower cooling rate) would reflect in a lower VT_Max_. Additionally, no difference in heating rate was found between species and only *P. nattereri* showed a significant difference in its VT_Max_ between day and night. Regarding habitat use, we found *P. cuvieri* to be slightly more abundant in open habitats than in non-open habitats, whereas *P. nattereri* shows the reverse pattern, which does not support our second prediction that the species with the lower thermal tolerance would not occur predominantly in open areas. Lastly, in spite of both species being widespread in Cerrado, they showed different patterns of VT_Max_ - ET_Max_ values throughout their ranges, with only *P. nattereri* having most of its records in localities with temperatures below its VT_Max_. Thus, only for *P. nattereri* did we confirm our prediction that global distribution is mostly comprised of localities with environmental temperatures below the VT_Max_.

The difference in VT_Max_ values between these two frog species might be related to their different body sizes [55–56] but additionally might reflect their natural history. Although there was no difference in heating rate between the species, *P. nattereri* might still cool slower when exposed to high temperatures. Since it burrows in the soil [33–34], this may allow it to quickly reduce its body temperature, since the soil is a good thermal insulant [57]. On the other hand, *P. cuvieri* uses preexisting cavities as diurnal refuges (e.g. see [58–59]), which, in spite of being below ground level, can heat up faster than the soil (S2 Fig). Yet, despite having a lower VT_Max_, most of the localities of *P. cuvieri* in Cerrado have temperatures above its VT_Max_. This suggests that other aspects of its thermal ecology might be playing a role in avoiding thermal stress, such as a reduced daily activity time or physiological traits regulating hydration state.

As wet skin ectotherms, hydration level can also influence the temperatures tolerated and selected by individuals for thermoregulation in their habitats [60–63]. This has been observed for other frog species (e. g. *Lithobates catesbeianus*; [15]), with individuals decreasing their VT_Max_ in response to dehydration, and some even losing their behavioral response to VT_Max_. Even though we controlled for hydration when measuring VT_Max_, individuals in the wild rarely are at their optimal hydration level and thus desiccation might influence local frog distribution [64]. Desiccation has been shown to be correlated with substrate use [65] and with dispersal probability throughout the landscape [64]. Additionally, closely-related frog species may vary in their response to desiccation along thermal gradients, with some species showing greater resistance to water loss at lower temperatures, and others at higher temperatures [66]. Therefore, knowing the interaction between VT_Max_ and hydration state of individuals in their environments can help to understand patterns and/or limits in their distribution [64, 67–69].

We found that *P. nattereri* was most abundant in non-open habitats, despite our second hypothesis. This may be related to the fluctuating daily temperatures of Cerrado savannas. While open areas reach higher temperatures (up to 35–37 °C in open habitats versus 32–35 °C in non-open habitats, *pers. obs.*) due to the exposed soil and low to no tree cover, they also cool rapidly in the evening. Non-open areas, on the other hand, do not get as hot but stay warmer for longer, which may expose individuals to a longer period with temperatures close to their thermal limits (e.g. [14]). This may favor the occurrence of *P. nattereri* in non-open habitats, since it can withstand higher temperatures before seeking refuge. Again, this may be related to evaporative water loss since desiccation is more intense in open habitats [70–71]. Even though we found a relatively high variation in the data on habitat use for both species, the difference in the use of open and non-open habitats between species seem to reflect the overall patterns of their distribution throughout the Cerrado. We highlight the importance of considering different spatial scales — geographic range and habitat use, as proposed by [72] because these allow to quantify how species distribution may reflect different aspects of their niches.

Despite numerous ecophysiological studies comparing how environmental temperatures influence habitat use of species [20, 73], these rarely account for thermal tolerances. Using behavioral thermal tolerances, such as the VT_Max_, allows for the integration of thermoregulatory behavior, which usually happens before critical limits are reached [3, 74–75]. Furthermore, combining the VT_Max_ with natural history and geographic distribution data can be critical to understand how future scenarios of global warming might impact distribution [76–77], especially for amphibians which are already under a global decline worldwide [78–79]. Our study indicates that differences in behavioral thermal tolerance may be important in shaping local and regional distribution patterns. Furthermore, small-scale habitat use might reveal a link between behavioral thermal tolerance and natural history strategies. Further studies using additional sympatric species of the genus *Physalaemus* (e. g. *P. centralis*, from the same clade of *P. cuvieri*, and *P. marmoratus*, from the same clade of *P. nattereri*) could help to elucidate if those differences are due to body size variation or if tolerances are phylogenetically conserved. We hope this study stimulates future mechanistic studies on amphibian thermal ecology and on the impact of global warming on species distribution.

## Acknowledgements

We thank Instituto Florestal for allowing our work at Estação Ecológica de Santa Bárbara (permit #260108-008.476/2014; ICMBio-SISBIO for the permit to collect frog specimens (permit #50658-3); and Fundação de Amparo à Pesquisa do Estado de São Paulo for a grant (#2018/14091-8). This study was financed in part by the Coordenação de Aperfeiçoamento de Pessoal de Nível Superior – Brasil (CAPES) – Finance Code 001”. MM thanks Conselho Nacional de Desenvolvimento Científico e Tecnológico for a research fellowship (# 306961/2015-6).

## Supporting information

**S1 Fig.** Detail of hind feet of (A-B) *P. nattereri* and (C-D) *P. cuvieri*, showing the inner and outer metatarsal tubercles in the detail. Note the much larger and strongly keratinized tubercles in *P. nattereri.* Photos not to scale.

**S2 Fig.** A) Temperature during a 24-hour cycle measured with sensors buried in the soil at superficial soil (green) and below ground level (red) and in a frog-sized plaster model (blue). B) Illustration of the measurement setup.

**S1 Table.** Voluntary Thermal Maximum measurements of both species.

**S2 Table.** Temperature data of *P. cuvieri* during experiments.

**S3 Table.** Temperature data of *P. nattereri* during experiments.

**S4 Table.** Geographical records of both species in the Cerrado ecoregion.

**S5 Table.** Habitat and abundance data for both species in six localities of the Cerrado ecoregion. Sources of data: Thomé (2006); Oliveira (2012); Duleba (2013); Ramalho et al. (2014); Motta (2019).

